# A method to estimate the contribution of regional genetic associations to complex traits from summary association statistics

**DOI:** 10.1101/024067

**Authors:** Guillaume Pare, Shihong Mao, Wei Q. Deng

**Affiliations:** Department of Pathology and Molecular Medicine, McMaster University, Hamilton, ON L8S 4L8, Canada; Population Genomics Program, Department of Clinical Epidemiology and Biostatistics, McMaster University, Hamilton, ON L8S 4L8, Canada; Population Health Research Institute, Hamilton Health Sciences and McMaster University, Hamilton, ON L8L 2X2, Canada; Thrombosis and Atherosclerosis Research Institute, Hamilton, ON L8L 2X2, Canada; Department of Statistical Sciences, University of Toronto, Toronto, ON M5S 3G3, Canada

## Abstract

Despite considerable efforts, known genetic associations only explain a small fraction of predicted heritability. Regional associations combine information from multiple contiguous genetic variants and can improve variance explained at established association loci. However, regional associations are not easily amenable to estimation using summary association statistics because of sensitivity to linkage disequilibrium (LD). We now propose a novel method to estimate phenotypic variance explained by regional associations using summary statistics while accounting for LD. Our method is asymptotically equivalent to multiple regression models when no interaction or haplotype effects are present. It has multiple applications, such as ranking of genetic regions according to variance explained or comparison of variance explained by to or more regions. Using height and BMI data from the Health Retirement Study (N=7,776), we show that most genetic variance lies in a small proportion of the genome and that previously identified linkage peaks have higher than expected regional variance.

## Introduction

Currently known genetic associations only explain a relatively small proportion of complex traits variance. In accordance with the widely accepted polygenic nature of complex traits, it has been proposed that weak, yet undetected, associations underlie complex trait heritability^1^. We have recently shown that the joint association of multiple weakly associated variants over large chromosomal regions contributes to complex traits variance^2^. Such regional associations are not easily amenable to estimation using summary-level association statistics because of sensitivity to linkage disequilibrium (LD). Nonetheless, only large meta-analyses have the necessary power to identify weakly associated variants. In this report, we propose a novel method to assess the contribution of regional associations to complex traits variance using summary association statistics. Estimation of regional associations can help identify key genomic regions involved regulation of complex traits.

Clustering of weak associations within defined chromosomal regions has been previously suggested^3^ and can increase variance explained at established association loci as compared to genome-wide significant SNPs alone^4^. Such regional associations extended up to 433.0 Kb from genome-wide significant SNPs^2^, a distance compatible with long-range *cis* regulation of gene expression^5^, ^6^. Furthermore, regional associations appeared to be the results of multiple weak associations rather than one or a few very significant univariate associations. These results point towards the existence of key regulatory regions where functional genetic variants aggregate, the identification of which can lead to novel biological insights and a better understanding of complex traits genetics.

Several methods have been described to estimate the overall contribution of common genetic variants to complex traits variance, but no method was specifically designed to estimate regional association using summary association statistics while accounting for LD. For instance, a popular approach is based on variance component models using genetic relatedness as the variance-covariance matrix of the random effect. An implementation of this approach uses REML^1^ to estimate the genetic effect, and variations have been reported to either take into account LD between SNPs^7^ or to handle large datasets. While very useful and informative, all of these approaches require individual-level data. This latter pitfall has been overcome by development of LDScore^9^, which uses summary-level association statistics as input and has been shown to be equivalent to Elston regression^10^. However, LDScore is ill suited for estimation of regional heritability because it requires regressing genetic effect as function of linkage disequilibrium over large number of SNPs. A multi-SNP locus-association method has been described that uses summary association statistics, but it requires to prune SNPs for LD (r^2^ < 0.1) first^11^. Finally, alternative methods^12^, ^13^ estimate genetic variance from the distribution of effect sizes and are not appropriate to estimate genetic variance from small regions, especially as linkage disequilibrium is expected to be strong in the case of regional associations. There is thus a need for a method to estimate the regional contribution of common genetic variants using summary association statistics and taking LD into account.

## Results

### Comparison of genetic variance estimated using summary statistics and variance component models

Our method estimates regional contribution to complex trait variance using summary data by adjusting the variance explained by each SNP for its LD with neighboring SNPs. As the method is equivalent to multiple regression models when no haplotype or interaction effects are present, we sought to compare estimation of overall genetic variance by our novel method to results of variance component models. We applied our method to BMI and height in the Health Retirement Study (HRS; *N* = 7,776) for which individual-level genotypes were available. We first divided the genome in SNP blocks minimizing inter-block LD and tested both methods, using only summary association statistics for our method and corresponding individual-level data for variance component models. Both methods provided consistent estimates of genome-wide genetic variance (Figure 1). We explored the impact of adjustment for genetic principal components. The first 20 components provided adequate protection against population stratification while inclusion of fewer components led to inflation in genetic variance, especially for height. Using 20 components, genetic variance was estimated at 0.12 (95% CI 0.01-0.24) for BMI with our novel method and 0.14 (95% CI 0.05-0.23) with variance component models^14^. Corresponding estimates for height were 0.28 (95% CI 0.17-0.39) and 0.30 (95% CI 0.20-0.39).

**Figure 1.**
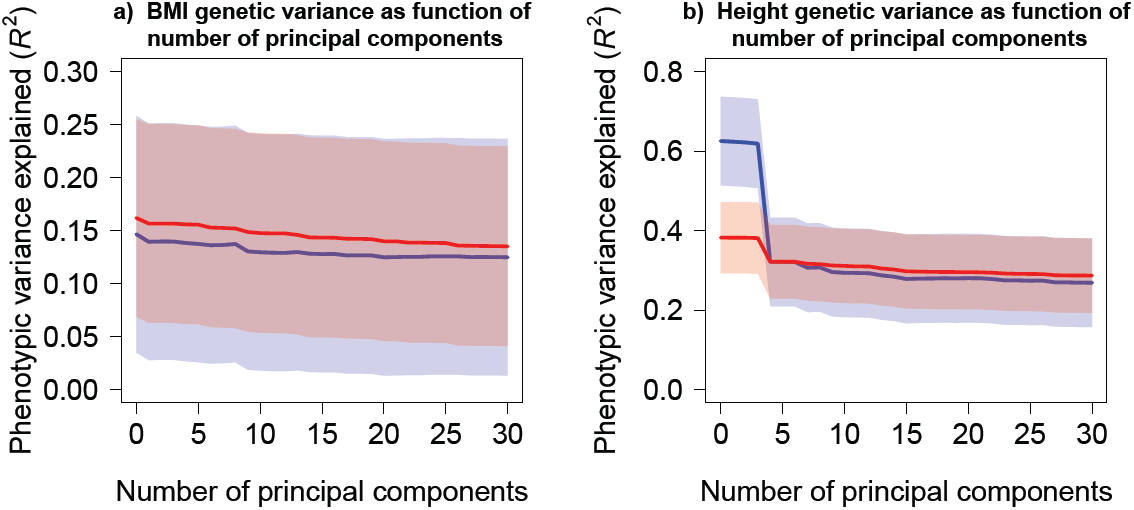
Genome-wide genetic variance estimated by summary association statistics and variance component models. The overall genetic variance estimated by summary association statistics and variance component models is illustrated as a function of number of principal components for BMI (Panel A) and height (Panel B). Blue lines represent estimates of genetic variance using summary association statistics; with 95% confidence intervals illustrated as blue shaded area. Corresponding estimates for variance component models are in red.

### Use of summary association statistics to identify regional associations

We next sought to compare the ability of |our method to identify regional associations with other approaches also using summary association statistics. We ranked SNP blocks (median size of 250 Kb) according to decreasing regional genetic variance by applying three different methods on GIANT summary association statistics: (1) our proposed approach, (2) LDScore and (3) the multi-SNP locus-association proposed by Ehret and colleagues. We then used SNP block ranks derived from GIANT using each of the three methods and estimated genetic variance in HRS (which was not part of GIANT) with variance component models, successively adding a higher proportion of HRS blocks. As illustrated in Figure 2, our novel approach provided better performance, with genetic variance increasing more rapidly as function of proportion top SNP blocks included. While the multi-SNP locus-association method performed well, a high proportion of SNP blocks didn’t have any SNP left after LD and *p*-value pruning (68.4% for BMI and 26.7% for height), highlighting the advantage of adjusting for LD in our method instead of LD pruning. We obtained consistent results when using 1000G data instead of HRS to derive the LD structure (Figure S2).

**Figure 2.**
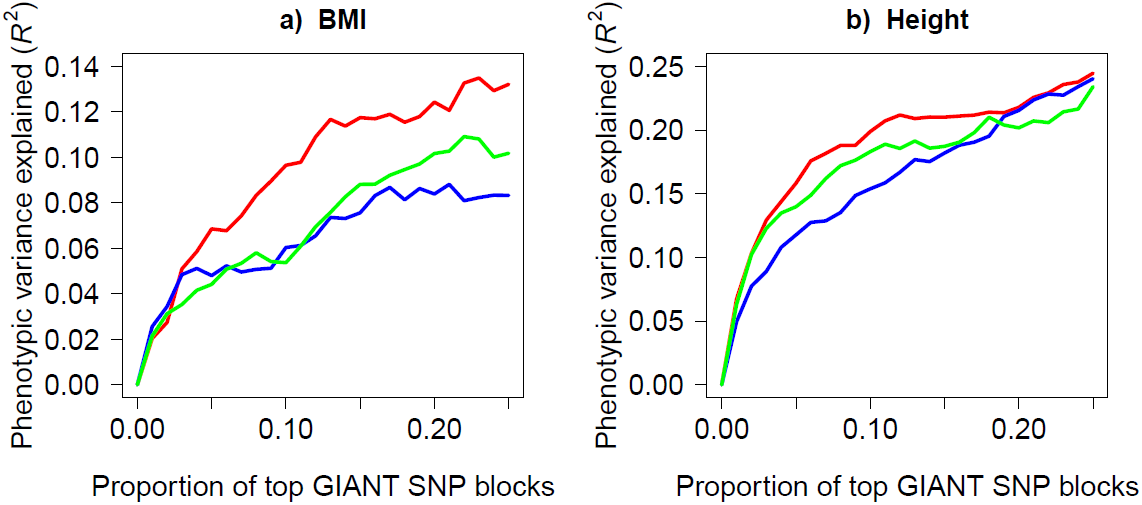
Genetic variance as a function of proportion of top SNP blocks. Genetic variance in HRS based on SNP block ranking derived from GIANT summary association statistics. Three methods were tested to rank SNP blocks: our novel approach (red), LD Score (blue) and multi-SNP locus-association (green). The median SNP block size was 250 Kb (i.e. 85-95 SNPs). Genetic variance was calculated in HRS using variance component models for BMI (Panel A) and height (Panel B).

A relatively small proportion of blocks contributed disproportionally to genetic variance. When dividing the genome into SNP blocks of median size 250Kb, corresponding to 85-95 contiguous SNPs, the top 25% SNP blocks explained 0.94 of BMI genetic variance (i.e. = 0.132/0.140) and 0.83 of height genetic variance using our method to rank SNP blocks. These results could potentially be explained by the presence of one or more very strong associations in each of these top SNP blocks. To explore this possibility, we recorded the minimum univariate association SNP *p*-value for each block in both GIANT and HRS (Figure 3). Median minimum univariate *p*-value was 2.0×10^2^for BMI in GIANT, with 0.01 of blocks having one or more genome-wide significant associations (*p* < 5×10^−8^). On the other hand, the median minimum *p*-value was 1.9×10^−3^for height in GIANT, with 0.09 of blocks having one or more genome-wide significant associations. The median minimum univariate *p*-values were 1.8×10^−2^ and 1.7×10^−2^ for BMI and height in HRS. No SNP reached genome-wide significance in HRS.

**Figure 3.**
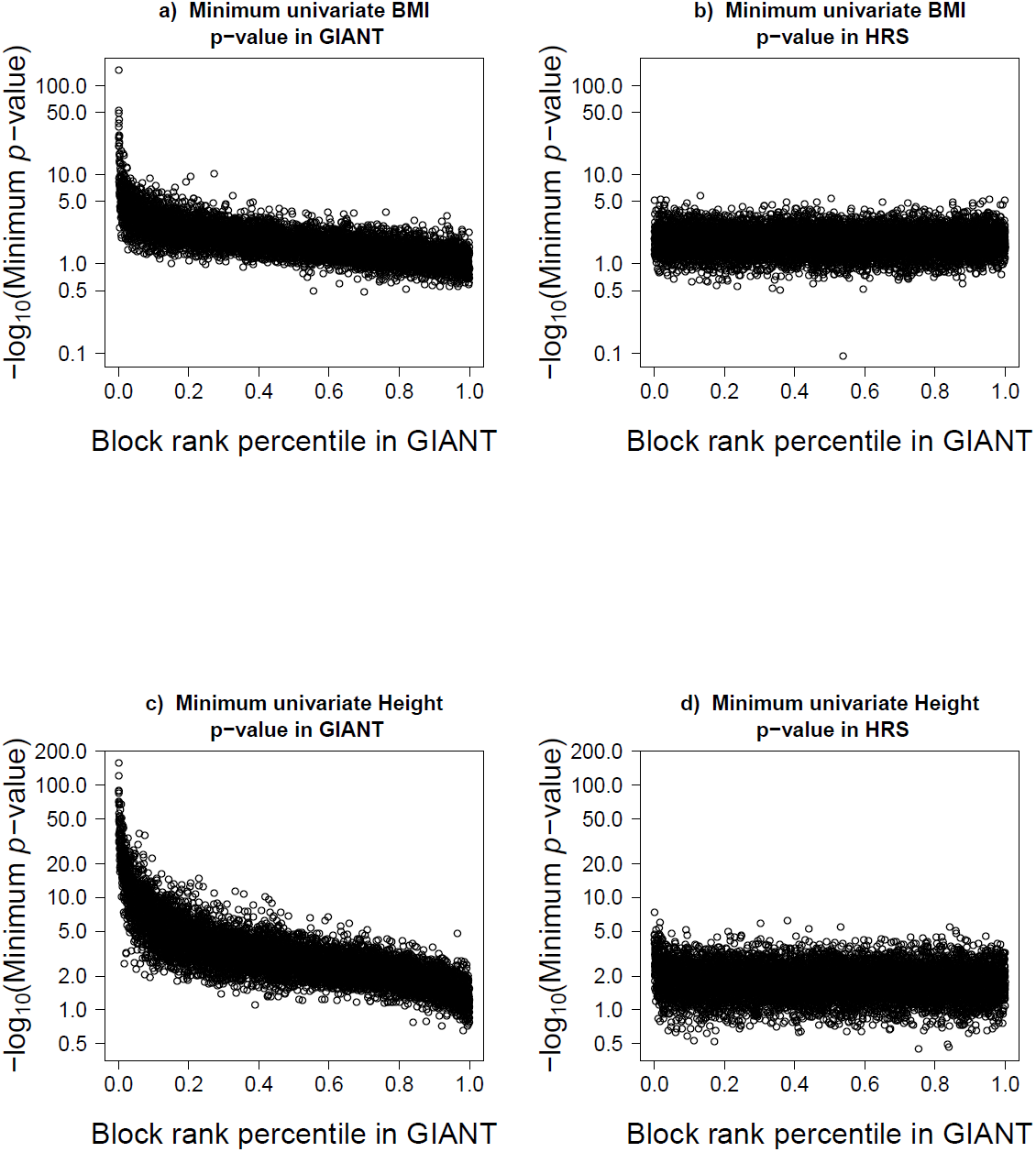
Minimum univariate SNP association p-value for each SNP block. SNP block ranks are based on GIANT data using our novel approach while the minimum univariate SNP association *p*-values were taken from GIANT (Panels A and C) or calculated in HRS (Panels B and D) for BMI (Panels A and B) and height (Panels C and D). The median SNP block size was 250 Kb (i.e. 85-95 SNPs).

### Analysis of known linkage peaks

A unique application of our method is the estimation of genetic variance over extended genomic regions using summary association statistics data. We therefore tested the hypothesis that previously identified linkage peaks are enriched for regional associations. Based on the largest single linkage study of height and BMI^15^, 3 peaks with suggestive (LOD>2.0) evidence of linkage in Europeans were identified, all for height. The only peak with LOD>3.0 showed a significant (*p* =0.002) enrichment in regional association within a distance of +/- 7.5 Mb of the linkage marker, corresponding to an estimated excess regional genetic variance of 0.0044 as compared to genome-wide average (Table 1). Upon closer inspection, the region encompasses several sub-regions with genome-wide significant associations.

**Table 1.** Excess regional genetic variance at 3 suggestive (LOD>2.0) linkage peaks for height using GIANT summary association statistics. Regional genetic variance was calculated within +/- 7.5 Mb of each peak marker and compared to genome-wide average for regions of equivalent size. P-values are two-sided.

| Chr. | Peak Marker | LOD Score | Excess genetic variance | 95% CI Upper limit | 95% CI lower limit | *p*-value |
| --- | --- | --- | --- | --- | --- | --- |
| 11 | D11S2000 | 2.74 | -0.0021 | 0.0002 | -0.0044 | 0.07 |
| 12 | D12S1301 | 2.07 | -0.0005 | 0.0014 | -0.0024 | 0.61 |
| 15 | D15S655 | 3.00 | 0.0044 | 0.0072 | 0.0017 | **0.002** |

## Discussion

We propose a novel method to estimate regional genetic variance from summary association statistics. Using this method, we confirmed a major role of regional associations in complex trait heritability, whereby the aggregation of genetic associations contributes disproportionally to phenotypic variance. Selecting the top SNP blocks from the GIANT meta-analysis, we showed that 25% of the genome is responsible for up to 0.94 and 0.83 of BMI and height genetic variance. A large proportion of these blocks had unremarkable minimum univariate *p*-values, suggesting the presence of multiple weak associations underlies their impact on phenotypic variance, especially for BMI. Concentration of genetic associations within these regions supports the existence of critical nodes in the genetic regulation of complex traits such as height and BMI, with implications not only for association testing but also for population genetics and natural selection. These results also suggest a combination of strong genetic associations and regional associations contribute to complex traits variance, with the relative proportions varying across traits. For instance, a higher proportion of genetic variance was found in the top 25% blocks for BMI yet these blocks had less significant minimum univariate *p-* value than height.

Our method can also be used to estimate the genetic variance explained by extended regions. We therefore tested the hypothesis some of the previously identified linkage peaks are the result of regional associations. The only peak with LOD>3.0 from the largest linkage study of height showed a marked and significant enrichment in regional association^15^. This region had been previously identified in other linkage studies^16^. Genetic variance explained by the region was estimated at 0.0044, which is unlikely to explain the linkage peak by itself. Nonetheless, the juxtaposition of linkage and regional associations points towards concentration of functional variants as a potential explanation for the observed linkage. The lack of regional association at other peaks can be explained by false-positive linkage results, the possibility rare variants not captured in GWAS studies underlie linkage peaks, or by differences in genetic architecture between studied populations.

Our proposed method has several advantages and is complementary to other methods. First, it provides results that are highly consistent with the widely used variance component models requiring individual-level data. Second, it is immune to “spillage” association caused by LD between SNP blocks as LD is accounted for, thus enabling blocks to be freely defined. Third, it is computationally straightforward and can be applied to large datasets. Fourth, it is agnostic and therefore complementary to functional annotations of the genome. Fifth, since SNPs are not pruned for either LD or significance, every SNP block or region can be evaluated.

A few limitations are worth mentioning. First, the assumption that SNPs contribute to genetic variance without any interaction or haplotype effects can lead to an underestimation of genetic variance. While our estimates were consistent with variance component models, there is a need for statistical models that more fully capture genetic variance, especially when strong haplotype effects are expected^17^. Second, estimation of overall genome-wide genetic variance using summary association statistics is dependent on both the accuracy of SNP effect size estimates and correct specification of LD structure. For instance, differences in adjustment for population stratification (e.g. principal components) in individual studies participating to a meta-analysis could potentially affect results. However, gross misspecification of LD is expected to bias results towards the null. Third, we have only tested continuous traits. Nevertheless, our method can be easily adapted to other outcome types through the use of generalized linear models.

In this report, we establish a novel method to estimate the regional contribution of common variants to complex traits variance using summary association statistics. Our method has many applications. By ranking genetic regions we showed regional enrichment in genetic associations for BMI and height. Identification of key genetic regions is also important for future fine-mapping studies. Indeed, our method can be used to perform network analysis using summary association statistics, or to combine summary association statistics with other types of genetic annotations such as linkage studies results, as we have done. Finally, our method can provide insights into patterns of genetic associations, such as the observed dissimilarities between BMI and height with respect to distribution of regional associations and response to regional gene scores.

## Material and Methods

### Methods overview

We have previously shown that large regions joint associations, where multiple genetic variants are included as independent variables in a linear model, are a simple and powerful way to test for regional associations when individual-level data are available^2^. However, such regional joint associations are not easily amenable to estimation using summary data because of sensitivity to linkage disequilibrium. To evaluate the contribution of large regions joint associations to variance of complex traits, we first devised an algorithm to divide the genome in blocks of SNPs in such a way as to minimize inter-block linkage disequilibrium and thus “spillage” associations. We then derived a method to estimate regional associations using summary data and showed this method to be equivalent to multiple regression models when genetic effects are strictly additive (i.e. no haplotype or interaction effect). Using regional variance estimates from summary-level association statistics for height and BMI from the Genetic Investigation of Anthropometric Traits (GIANT) consortium, we estimated the distribution of genetic effects across the genome and validated results in the Health Retirement Study (HRS), which was not part of GIANT.

### Dividing the genome into SNP blocks

We first divided the genome into regions of contiguous SNPs varying in size (e.g. from 195 SNPs to 205 SNPs), herein referred as SNP blocks and used as units for regional associations. To minimize inter-block LD and thus “spillage” associations, we devised a greedy algorithm optimizing choice of block boundary sequentially from one end of a chromosome to the other. Briefly, using user-defined minimum and maximum block size (in number of SNPs) and starting at one end of a chromosome arm, each possible “cut-point” between the first and second block are tested and maximal LD (*r^2^*) between pairs of SNPs crossing block boundary is calculated. The cut-point that minimizes maximal LD is chosen, thus defining the first block, and the procedure is repeated for each subsequent block until all SNPs on a chromosome arm have been assigned to a block. We empirically determined that SNP blocks of size 85 SNPs to 95 SNPs (median 90 SNPs) had a median physical size of 250 Kb.

### Estimating the contribution of regional genetic associations with individual-level genotypes

Use of adjusted *R^2^* lends itself nicely to estimation of regional variance explained when individual-level genotypes are available^2^. In this context, SNPs comprised in a given SNP block are included as independent variables in a multiple linear regression model and the goodness of fit statistic, adjusted *R^2^* calculated. Because adjusted *R^2^* accounts for the number of SNPs included in each block, expected adjusted *R^2^* is zero under the null of no association and the expected sum of adjusted *R^2^* over all SNP blocks is also zero. The overall contribution of regional associations to complex traits variance can be estimated by simply summing adjusted *R^2^* over all (or selected) SNP blocks.Furthermore, the distribution of adjusted *R^2^* under null has been previously described^18^ and can be used to derive the distribution of the sum of adjusted *R^2^*.

### Estimating the contribution of regional genetic associations with summary association statistics

Estimating the contribution of regional genetic associations from summary association statistics is challenging when the exact SNP linkage disequilibrium structure of source populations is unknown. While approaches have been described to perform joint or conditional associations^17^, ^19^ using estimated SNP covariance matrices, they do not perform well when estimating regional variance explained because of sensitivity to misspecification of linkage disequilibrium (data not shown) and ensuing overestimation of regional associations. We therefore created a simple procedure to estimate regional variance explained from summary association statistics data, adjusting for linkage disequilibrium.

Without loss of generalizability, we assume a quantitative trait (*Y*) standard normally distributed and genotypes normalized to have mean = 0 and standard deviation = 1 throughout. Given an *n* x *m* genotype matrix *X* representing genotypes at *m* SNPs in *n* individuals and the pairwise linkage disequilibrium (*r^2^*) between two SNPs *k* and *l* as *r^2^ _k,1_*, for a SNP *d*, the following LD adjustment (*η_d_*) can be defined as the summation of LD between the *d^th^* SNP and 100 SNPs upstream and downstream:

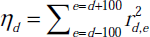

with a distance of 100 SNPs assumed sufficient to ensure linkage equilibrium (other values might be used). Only including SNPs with summary GWAS statistics in the sum, variance explained by each SNP *d* is given by:

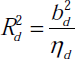

where *b_d_* denotes the univariate regression coefficient commonly reported in GWAS results. Regional variance explained is then given by the sum of 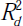 over SNPs comprised in a given region. Assuming a strictly additive genetic model where each SNP contributes additively to a trait without any interaction or haplotype effects, we demonstrate the expected total variance explained over a region 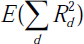, is approximately equal to the expected value of the multiple linear regression variance explained *E(R^2^)* when the sample size is sufficiently large.

To simplify the calculation, we define *D* such that 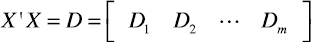 is an *m* by *m* symmetric matrix, where *D_k_* is a *m* x *1* vector whose entries represent the *k^th^* column of *D*. We will make use of the following properties:

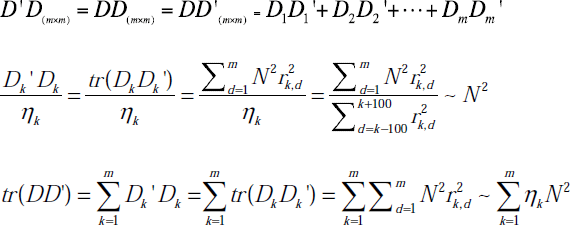

#### Estimation of regional genetic variance with multiple linear regression models

Suppose the genotype matrix is fixed while the true genetic effect is a random vector β, whose individual components, i.e. the SNPs, *i* = 1,2,···,*m*, have mean 0 and variance *σ^2^*. The size of the variance *σ^2^* is on the scale of 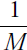, where *M* is the number of genome *M* wide SNPs. The genetic model can be expressed as:

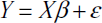

where *ε* is a vector of standard normal error with identity variance covariance matrix. Then, the vector of estimated multiple linear regression coefficients *B* is given by:

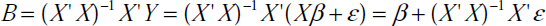

The multivariate variance explained *R^2^* can be written in terms of the true effect and the error term:

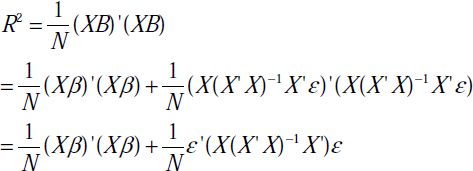

and since the error term has identity variance covariance and the true effect *β* has variance covariance matrix *σ^2^1*, the expected variance explained is simplified to

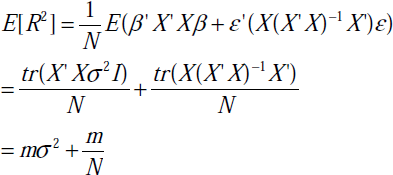

and the variance can be calculated accordingly:

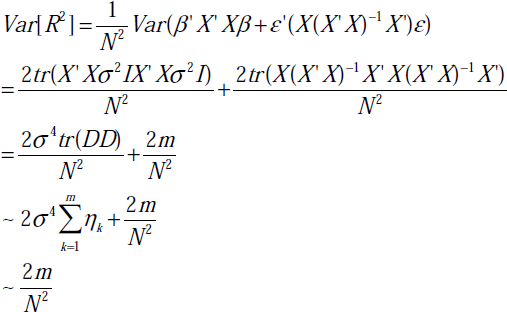

#### Estimation of regional genetic variance using summary association statistics

The univariate regression coefficients, denoted| by lower case *b* and directly obtained from GWAS summary statistics, are given by

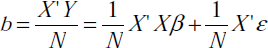

The total variance explained over a region 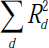 can be calculated using only the univariate regression coefficients from GWAS:

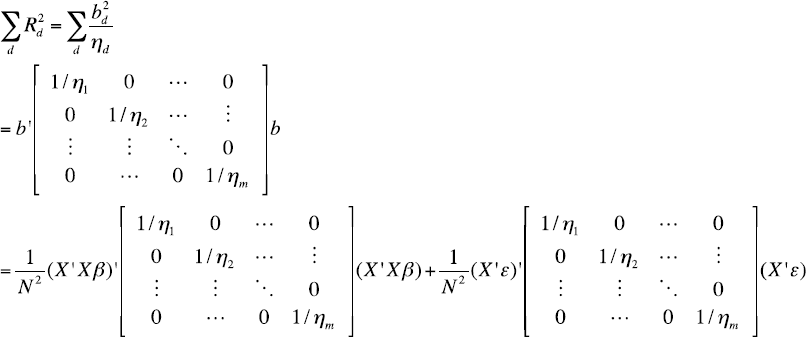

We can rewrite the expression by defining:

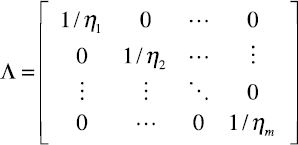

Thus, we have:

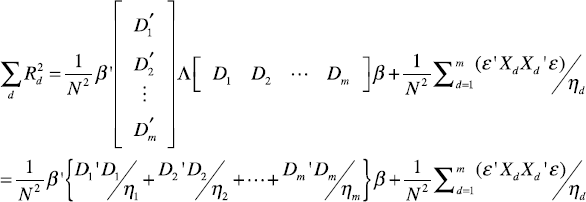

Since 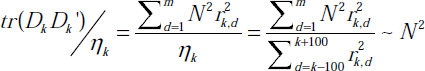 as LD tends to be weak 100 SNPs upstream and downstream away from the index SNP, we can simplify the expected total variance explained using the summary statistics to:

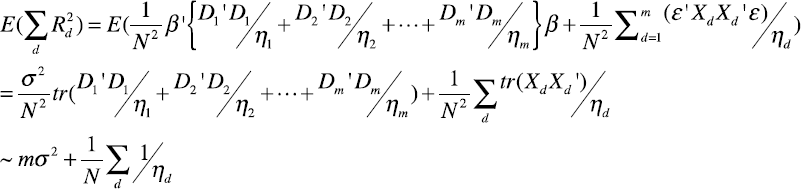

The variance of 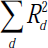 can be similarly derived. By using the cyclic property of trace, and the fact that *DΛD* is a positive definite matrix, an upper bound of the variance can be expressed as:

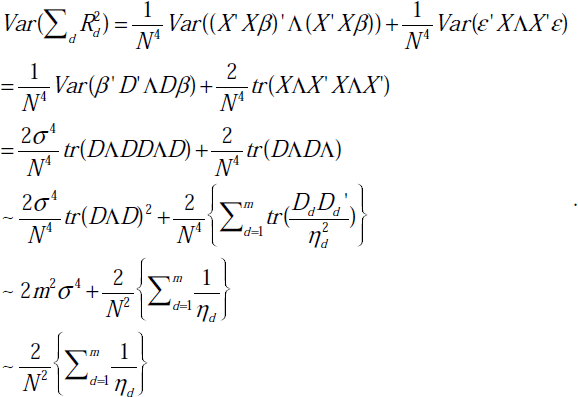

Where 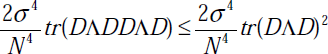 and both terms are small with respect to 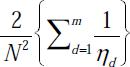.

#### Comparison of regional genetic variance estimated using multiple regression models and summary association statistics

The expected values are equivalent between multiple linear regression 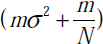 and the summary statistics 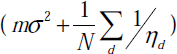 derived regional sum, with the number of SNPs *m* replaced by the “effective” number 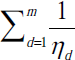 Variance is slightly bigger for multiple linear regression models 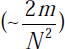 as compared to regional sum 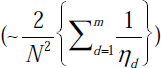 because the “effective” number of genetic markers 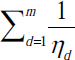 is always equal (no LD) or less than the number of markers (*m*). In other words, *Var* 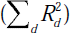 is expected to be equal (no LD) or lower than the corresponding *Var* (*R^2^*).

### Health Retirement Study

We conducted large region joint association analysis for height using genome-wide data from the publicly available Health Retirement Study (HRS; dbGaP Study Accession: phs000428.v1.p1). HRS quality control criteria were used for filtering of both genotype and phenotype data, namely: (1) SNPs and individuals with missingness higher than 2% were excluded, (2) related individuals were excluded, (3) only participants with self-reported European ancestry genetically confirmed by principal component analysis were included, (4) SNPs with Hardy-Weinberg equilibrium *p* < 1x10^−6^ were excluded, (5) individuals for whom the reported sex does not match their genetic sex were excluded, (6) SNPs with minor allele frequency lower than 0.02 were removed. The final dataset included 7,776 European participants genotyped for 740,748 SNPs. Height and BMI was adjusted for age and sex in all analyses. To mitigate the effect of outliers, we have removed values outside the 1^st^ and 99^th^ percentile range for each of height and BMI. All analyses are adjusted for the first 20 genetic principal components unless stated otherwise. All LD estimates used throughout the manuscript were derived from HRS genotypes. HRS was not part of the GIANT meta-analysis of height and BMI^20^,^21^.

